# Color Desaturation in the Periphery is Explained by General Mechanisms of Contrast Sensitivity and Constancy

**DOI:** 10.1101/2025.02.28.640610

**Authors:** Ana Rozman, Jasna Martinovic

## Abstract

Colors appear less saturated in the visual periphery than in the fovea. We revisit this well-known phenomenon by characterizing parafoveal perceived contrast as a function of size. Observers (n = 20) matched perceived contrast of a parafoveally presented comparison disc (2° -0.33°) to a standard 2° disc. For chromatic stimuli, desaturation increased with decreasing size. Unexpectedly, a similar amount of desaturation occurred for luminance-defined discs, once their perceived contrast was adjusted to match the standard chromatic discs. Desaturation was reduced as standard stimulus contrast increased, in line with contrast constancy theory, which predicts constant appearance for stimuli that are sufficiently distant from threshold. Since chromatic contrast sensitivity is reduced away from the fovea, contrast constancy is unachievable within the monitor gamut. In conclusion and somewhat counter-intuitively, the appearance of color and luminance in the periphery is affected similarly, governed by general laws of contrast sensitivity and constancy.

**Public Significance Statement:** In the early stages of visual signal processing, color and luminance (achromatic) information are separated. If we overgeneralize on the basis of such early separation, we may conclude that the desaturated appearance of small and peripherally presented color stimuli is specific to color processing mechanisms. Here we show that this is not the case and that chromatic and achromatic stimuli are processed similarly when presented in the periphery, implying that such processing is governed by more general contrast-processing mechanisms. Our work provides the basis for a better understanding of how the visual system builds a unified experience of color appearance across chromatic and achromatic dimensions from the information on contrast present in the image.

## Introduction

Saturation is a perceptual property of color that describes the vividness of its hue. Saturation depends not only on a stimulus’ colorfulness (i.e., chromatic content) but also on its brightness (i.e., achromatic content; see Fairchild, 2013). For example, adding white light to a single-wavelength light would desaturate it. Thus, saturation increases with an increase in colorfulness but also decreases with an increase in brightness/lightness. An important distinction between brightness and lightness is that lightness is relational – by normalizing brightness relative to a surface that would be perceived as white, it can account for changes in illumination. However, saturation is not relational – it can also be experienced from isolated stimuli (e.g., a traffic light seen at night; Fairchild, 2013).

A more general concept than saturation is perceived contrast – referring to the overall amount of contrast perceived within a stimulus, irrespective of whether it is defined by color or luminance. Perceived contrast varies depending on the stimulus’ spatial properties and location on the retina. Chromatic patches, in particular, appear desaturated when presented parafoveally (McKeefry et al., 2007). Furthermore, chromatic patches of smaller sizes appear desaturated compared to patches of larger sizes (Abramov et al., 1991; Knau & Werner, 2002). These well documented phenomena can be linked to clear demonstrations of saturation constancy in the periphery -desaturation of peripheral stimuli is not experienced as long as their size is increased to account for the cortical magnification factor (Tyler, 2015). However, loss of perceived contrast (i.e., desaturation) in periphery does not need to be color-specific -if stimulus appearance is accounted for by general contrast constancy models, changes in the periphery should be similar across color and luminance, once they are matched in perceived contrast.

Suprathreshold appearance depends on contrast sensitivity. Contrast sensitivity function (CSF) captures how detection thresholds relate to spatial properties of the stimulus. Sensitivity is the inverse of threshold, i.e., the lower the amount of contrast required at a given spatial frequency for the stimulus to be detected, the higher the sensitivity for this frequency. For luminance-defined stimuli, this follows a bandpass curve (Campbell & Robson, 1968). Contrast sensitivity reduces due to multiple factors, for example with eccentricity (Rovamo et al., 1978), while higher spatial frequencies are specifically affected by ageing (Ashraf et al., 2020; Ross et al., 1985). Contrast sensitivity function can also be measured for chromatic patches. Chromatic CSF follows a low-pass shape for the yellow-blue and red-green contrast alike, with sensitivity dropping off faster with an increase in spatial frequency for yellow-blue contrast, compared to red-green (Mullen, 1985a). Chromatic contrast sensitivity is particularly reduced in the periphery (Gordon & Abramov, 1977; Johnson, 1986; Mullen, 1991). In fact, sensitivity for chromatic stimuli shows a notable drop at the upper end of the curve (i.e. at higher frequencies) with increasing parafoveal presentation (Ashraf et al., 2024). This is in line with Tyler’s (2015) demonstration of how retinal eccentricity can be compensated for by an increase in stimulus size, as reductions in stimulus size limit the range of spatial frequency detectors involved in processing the stimulus.

The specificities of vision at or near detection threshold cannot be fully interpolated to suprathreshold vision (Cannon, 1985; Vanston & Crognale, 2018). Still, the domains of threshold and suprathreshold vision are linked through the notion of contrast constancy (Georgeson & Sullivan, 1975), which describes perceived contrast of a suprathreshold stimulus as a function of contrast detection threshold for the same stimulus. This theory was initially defined for achromatic, luminance defined stimuli, but could also explain peripheral color desaturation. The higher the detection threshold for a certain spatial pattern, the higher suprathreshold contrast required to make it equal in perceived contrast to a different spatial pattern associated with a lower detection threshold. Since detection thresholds increase with eccentricity from the fovea for achromatic and chromatic stimuli alike, the decrease in saturation could be described as a consequence of contrast constancy– i.e., higher absolute contrast is required to rectify the loss of sensitivity in the periphery. Translated to stimulus size, increasing the stimulus as it moves towards the periphery rectifies the loss of sensitivity in higher spatial frequency channels. Therefore, rather than being reflective of color-specific low-level processing constraints, peripheral desaturation phenomena could rather be a governed by the same general mechanisms of contrast constancy, which have been well-described in relation to achromatic contrast processing.

In this study, we characterize desaturation as a function of size away from the fovea for colors that isolate the two cone-opponent mechanisms and compare it between chromatic and luminance isolating stimuli matched in perceived contrast. We also evaluate desaturation following reduction in stimulus size for chromatic stimuli presented on pedestals of varying levels of luminance contrast, using stimuli with a sharp and a blurred edge. Combined, our experiments will answer the following question: is desaturation in the periphery specific to color or is it governed by more general contrast constancy principles that apply equally to luminance-defined stimuli? If contrast constancy generalizes across chromatic and luminance contrast, we would expect to observe a similar desaturation with size across chromatic and luminance-defined stimuli. On the contrary, if desaturation with size in the periphery is a color-specific phenomenon, we should fail to observe a reduction in perceived contrast for luminance-defined stimuli. In our experiments, participants match perceived contrast of a comparison stimulus to that of a standard, fixed-size stimulus. If desaturation away from the fovea with a reduction of stimulus size is fully explained by contrast constancy, we should observe a gradual reduction of desaturation for both chromatic and achromatic stimuli as standard stimulus contrast increases and moves away from threshold. Additionally, if desaturation in the parafovea always occurs for color, addition of luminance contrast pedestals should not markedly reduce it, regardless of the luminance contrast added. On the contrary, if desaturation is a product of general principles that apply equally to luminance and chromatic vision, then it should follow a pattern predicted from these models when luminance pedestals are added. As a final test for this, we directly manipulate the edge of the standard disc to evaluate the dependence of perceived contrast on information carried by lower and higher spatial frequency channels.

## Methods

### Participants

A total of 20 participants (15 female, aged 20 – 32 years, M_age_ = 26) took part in the experiments. No participants were excluded. Sample size was verified with an a-priori power calculation in G*Power (Faul et al., 2009), based on data from comparable work (Abramov et al., 1991). We extracted their data in condition most comparable to our stimuli (4 color directions at 5 degree eccentricity and 0.25 – 2 degrees of visual angle in size) to calculate their observed power of the desaturation with size effect for color stimuli. This was the weakest for their ‘blue’ condition (Cohen’s *d* estimate = 1.10, ∼16% desaturation) with their sample size of 6. In order to replicate this with power of 0.8, we would require a sample of 15 participants. For their strongest effects (‘green’ and ‘yellow’ conditions, ∼54/58% desaturation) we would require a sample size of 5. Sufficiency of our sample size has been confirmed with post-hoc power simulation-based analyses (Kumle et al., 2021), recommended for linear mixed effect models, where it is more difficult to estimate all the relevant values a-priori (see Supplementary Materials for a detailed overview). This analysis verified that at least 0.8 power for all effects in all experiments would be achieved with a sample of 8 participants.

All observers had normal color vision as checked by City University Colour Vision Test (Fletcher, 1984) and normal or corrected to normal visual acuity according to an EDTRS chart check for viewing at 68 cm distance. Written informed consent was obtained from participants. The study was conducted in line with the Declaration of Helsinki (1964) and received ethical approval from University of Aberdeen School of Psychology Ethics Committee.

### Apparatus

Stimulus presentation was controlled by a VISaGe visual stimulus generator (Cambridge Research Systems, UK) and presented on a Display++ monitor (CRS, UK), using CRS toolbox and CRS color toolbox (Westland, Ripamonti, & Cheung, 2012) for MATLAB (Mathworks, USA). Responses were collected using a Cedrus RB-540 button box (Cedrus, USA). Colors were calculated on the basis of monitor spectra measurements (SpectroCAL, CRS, UK) and 2° cone fundamentals (Stockman & Sharpe, 2000). Participants were seated in a dark room at a 70 cm distance from the display, which was the only source of light.

### Stimuli and Procedure

Colors used in the experiment were defined to isolate the cardinal directions of chromatic mechanisms as represented in the DKL color space (Derrington, Krauskopf, & Lennie, 1984). Background was set to 0.3127, 0.3290, 22.9298 cd/m2 in CIE 1931 xyY coordinates, which is metameric to the neutral illuminant D65.

## Baseline Measurements

To ensure the absence of luminance contrast from color-only stimuli, we measured the point of isoluminance for both red-green and yellow-blue colors for each individual observer (see Figure 1 for an outline of experimental steps). We used a heterochromatic flicker spectrometry (HCFP, Walsh, 1958) task for this purpose. Observers were presented with two 2° circles flickering parafoveally, 5.5° to the left and right from central fixation. These corresponded to the standard stimulus size and location in the main experimental contrast adjustment task. The flicker was presented at the frequency of 20 Hz. Each of the presented patches flickered between two colors, red-green or blue-yellow, corresponding to L-M and S-(L+M) mechanisms. Observers were asked to minimize the perceived flicker between the two colors whilst fixating on the center of the screen. This was done by adjusting their luminances using a button box. Observers’ responses manipulated the angles of elevation in DKL space (Figure 1) in opposite directions – a left button press would increase the angle for color 1 and decrease it for color 2, and vice versa. The step size was set at a 4° luminance elevation for the RG and 1° for BY trials, corresponding to ±0.02 and ±0.12 RMS contrast units for the RG and BY mechanisms, respectively. The initial difference between the two colors in a trial was set between 1 and 5 multiples of the step size for that mechanism. Once the flicker was perceptually minimized, i.e., the point of isoluminance for the two colors was reached, observers pressed the top button to finalize their setting and proceeded to the subsequent trial. A total of 8 trials per color direction was performed. The individual isoluminance was obtained by calculating the mean luminance adjustment value for each of the color pairs based on 6 measurements, following the exclusion of the highest and lowest adjustment as outliers. The task took about 10 minutes to complete.

**Figure 1.**
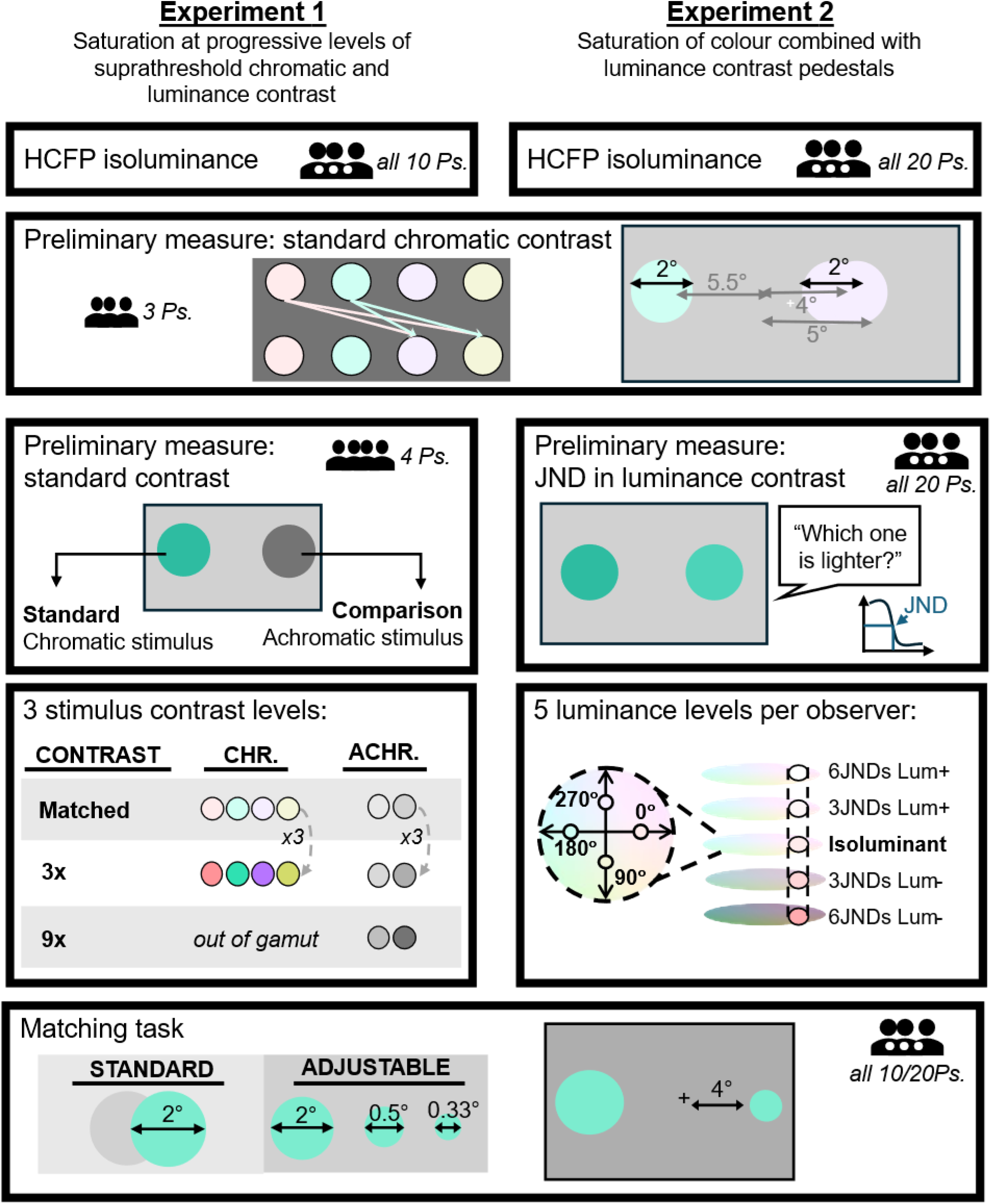
Schematic Representation of the Experimental Paradigm. Both experiments started with participants completing a HCFP task for establishing individual isoluminance (Panel 1). A subgroup of 3 participants completed an asymmetric matching task between cardinal colors of DKL color space (Panel 2). The standard contrast levels obtain with this baseline measurement were then used to match them to achromatic stimuli by a subgroup of 4 participants (Experiment 1), and to obtain JNDs in luminance contrast for all participants for Experiment 2 (Panel 3). The main experimental task was the same saturation/contrast adjustment task in both experiments (Panel 4). Standard stimulus, always 2° in size was presented parafoveally at 5.5° distance from fixation. The edge of standard stimulus varied between conditions and was sharp of blurred. The adjustable stimulus varied in size between conditions and was presented at 4° eccentricity. In the adjustment task, the standard and adjustable stimulus were always presented in the same color or luminance polarity and at the same level of contrast. The position of the stimuli (left or right of the fixation cross) was counterbalanced between conditions for all experiments.

To ensure that standard stimuli defined along different chromatic and luminance axes were equally salient to each other, perceived contrast was matched between 2° parafoveal discs in two baseline adjustment tasks (Switkes & Crognale, 1999). Firstly, three observers (one of the authors) completed an asymmetric matching task comparing red and green standard discs to yellow and blue comparison discs (Figure 1). This was done to ensure equal perceived contrast of standard stimuli between the cardinal colors of the two cone-opponent mechanisms. In this task, discs with a 2° diameter were presented at 4° to the left and right of the central fixation cross. The standard was either red or green, set at 0.02 RMS contrast. The comparison stimulus was either yellow or blue. Observers were asked to adjust the contrast of the comparison stimulus to match that of the standard stimulus whilst fixating on the central cross (i.e., using their peripheral vision). Stimuli were set to individual isoluminance, as measured with HCFP. Each standard and comparison stimulus pair were shown 8 times. The location of the stimuli in relation to the central fixation cross was counterbalanced. Results are visualized in Figure 2a.

**Figure 2.**
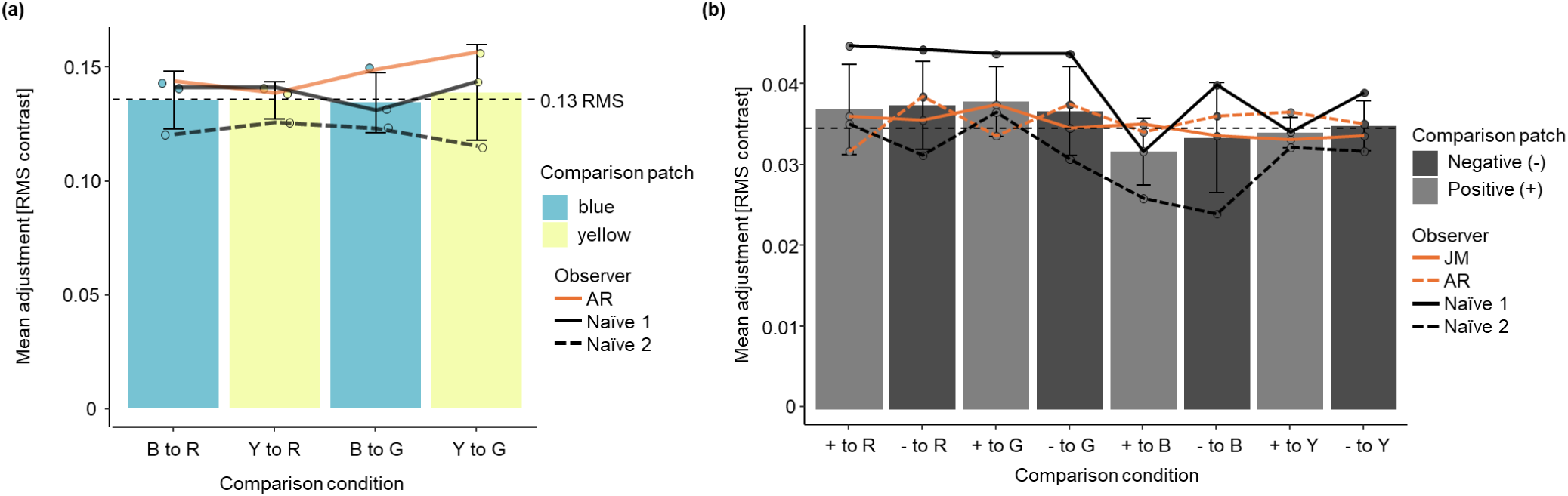
Results of Asymmetric Contrast Matching Baseline Tasks. Adjustments in baseline asymmetric matching task for (a) chromatic and (b) luminance stimuli. (a) Standard L-M stimuli were set to 0.02 RMS contrast. Error bars denote +/- 1 standard deviation between observers. The horizontal line corresponds to 0.13 RMS contrast which was used for standard stimulus contrast for the blue-yellow mechanism in the main experiment. Matches made by one author participant are marked with an orange line. On the x axis R, G, B and Y stand for reddish, greenish, bluish and yellowish, e.g. ‘B to R’ denotes bluish matched to a reddish standard. (b) Mean contrast adjustment is provided for positive (denoted by + on the x axis) and negative (denoted by – on the x axis) luminance polarity stimuli matched to chromatic standards in cardinal colors of the cone-opponent DKL color space (R, G, B and Y are used to denote these in the same way as in panel a). Error bars show the standard deviation between observers. A horizontal dashed line denotes the matched RMS contrast value used for luminance contrast stimuli (0.036). Matches made by authors are marked in orange.

Secondly, four observers (two of them authors) completed an asymmetric adjustment task between achromatic and chromatic stimuli. Two discs 2° in diameter were presented 4° to the left or right of the central fixation cross. The standard stimulus was always presented in one of the four cardinal colors in the cone-opponent color space (red, green, blue or yellow). The standard was set to individual observer isoluminance measured with HCFP. The contrast of the standard was set to the value established in the chromatic perceived contrast baseline measurement (see description of asymmetric chromatic contrast matching above). The comparison stimulus was of either positive or negative luminance polarity, set in the DKL space by fixing the angle of elevation to 90° (positive) or –90° (negative). Whilst fixating the central cross, observers were asked to adjust the contrast of the achromatic comparison so that it appears equally salient as the chromatic standard (Figure 1). Each measurement was performed 8 times, with the location of the standard counterbalanced. The grand-mean match was 0.036 RMS contrast (Figure 2b). There was no time limit for making a match. Observers completed 64 trials (4 colors, 2 polarities, 8 trials per condition) in about 20 minutes.

An additional baseline task was completed for Experiment 2, to create individually meaningful luminance contrast pedestals for the saturation adjustment task. A just noticeable difference (JND) in luminance contrast was measured for each observer (Figure 1, panel 3, right). This was presented as a 2AFC task staircase, asking observers to indicate which of the two presented stimuli is lighter. The two circles appeared on the screen briefly for 200 ms to prevent eye movements towards the stimuli, since a brief stimulus with luminance contrast could be expected to attract eye movements in an exogenous fashion (for literature on the timing of saccadic eye-movements see Hutton, 2008; Pierce et al., 2019; Salthouse & Ellis, 1980). Furthermore, observers were instructed to fixate the cross and avoid eye movements throughout the task.

Conditions were blocked by stimulus size and blocks were presented in a random order. The stimuli were either 2°, 0.5° or 0.33° discs and were presented at 4° eccentricity. Each block contained 8 interleaved staircases -two for each of the four cardinal colours, measuring either a positive or a negative luminance contrast JND. Each block ran until the termination criterion of 10 reversals was met for all of the 8 staircase procedures with Weibull function, implemented using the Palamedes toolbox (Prins & Kingdom, 2018). Approximately 50 matches on average were required for convergence of a single staircase. Observers completed the task in approximately 60-90 minutes. A screen encouraging observers to take a break was presented after each set of 80 completed trials. Before commencing each size block of the task, observers completed a short training block to get them adjusted to the task and stimulus size. For the first trial of each block, the luminance difference between the stimuli was set to a relatively high value of 1 cd/m^2^ to ensure the observers could locate the lighter stimulus comfortably and allocate their attention to the locations in which the stimuli would appear. This trial was excluded from the analysis.

### Experiment 1: Saturation at progressive levels of suprathreshold chromatic and luminance contrast

To compare the effect of desaturation in the periphery, we collected contrast adjustment data for isoluminant chromatic stimuli and luminance-only stimuli at multiple values of standard contrast. The task was completed by 10 of the total 20 observers (8 female, aged 20 – 32 years, M_age_ = 26).

Luminance standards were presented at three levels of contrast. This was either perceptually matched to contrast of chromatic isoluminant standard stimuli (as determined in the baseline measurements described above) or set at 3- or 9-fold multiples of that standard contrast. Color standards could only be presented at a 3-fold increase in contrast, as it was the maximum that could be achieved within the gamut of the monitor. The standard and comparison stimuli always appeared in the same color or polarity. The stimuli were presented at 4° eccentricity from the central fixation cross, either to the left or to the right, with the location counterbalanced. The standard was 2° while the size of comparison was either 2°, 0.5° or 0.33° of visual angle. Participants adjusted the contrast of the comparison stimulus using a button box, with the left button increasing the contrast and the right button decreasing the contrast, until a perceptual match was achieved. Participants were asked to complete the task whilst fixating a centrally positioned cross. Each of the 36 conditions (3 levels of contrast, 2 polarities, 3 sizes and 2 standard edge conditions) for the achromatic matching was presented 8 times, for a total of 288 trials. There was a total of 256 matches made for the chromatic matching task (32 conditions -2 levels of contrast, 4 colors, 3 sizes; presented 8 times each).

### Experiment 2: Saturation of lighter and darker colored discs with sharp and blurred edges

In the second part of the study, all 20 observers completed the same adjustment task but with chromatic contrast presented either at isoluminance or upon progressive luminance pedestals (i.e., lighter or darker than the background).

First, to match the amount of contrast between observers, Just Noticeable Differences (JNDs) for discriminating positive and negative luminance polarity contrasts combined with chromatic contrast (red, green, blue and yellow) were measured for each participant (details in Baseline Measurements section).

Colors to be used for each observer were based on these baseline matches. In addition to an isoluminant chromatic setting (identical to the one used in the first experiment), the same chromatic information was also presented with an addition of a positive or negative luminance pedestal, equal to a value of 3 or 6 multiples of JNDs for a given color direction. In the main experimental task, participants were again asked to adjust saturation of a comparison stimulus to match that of a standard stimulus whilst fixating the central cross. The two circular stimuli were presented at 4° eccentricity left or right of the foveal fixation cross and always appeared in the same color (reddish, greenish, bluish or yellowish) and luminance pedestal (isoluminant, 3 or 6 JNDs positive or negative luminance contrast pedestal). The location of the standard and comparison stimulus was counterbalanced, and the presentation of colors randomized. All conditions with the same luminance contrast pedestal were blocked (split into two parts to allow for breaks) and the order of these pedestal blocks was randomized.

The standard stimulus was always a 2° disc whilst the size of the comparison stimulus was either 2° (baseline comparison), 0.5° or 0.33° of visual angle. Observers were asked to adjust the amount of chromatic contrast of the comparison stimulus to match the saturation of the standard stimulus using a button box (left button increasing, right button decreasing). Adjustments were made 8 times per condition (4 colors, 3 comparison stimulus sizes, 5 luminance contrast pedestal conditions and two standard stimulus edge definitions; see Fig. 1), with a total of 960 trials. Trials for two standard stimulus edge conditions were split into two testing sessions, each lasting up to 2 hours for participants to complete, depending on chosen duration of breaks.

Finally, this experiment also manipulated stimulus spatial properties by blurring the edge of the standard disc to reduce the information carried by the higher spatial frequency channels. The stimulus edge was blurred by adding an outer ramp encompassing 15% of the stimulus area, within which contrast was reduced linearly (Figure 3a). This resulted in 7.5% difference in overall contrast between the sharp and blurred edge standard stimulus. The standard 2° stimuli (sharp edge and blurred edge) still have comparable amount of information in the lower spatial frequency range, but the blurred edge standard stimulus is better matched in energy to smaller, sharp edge comparison stimuli in the higher frequency domains (Figure 3b).

**Figure 3.**
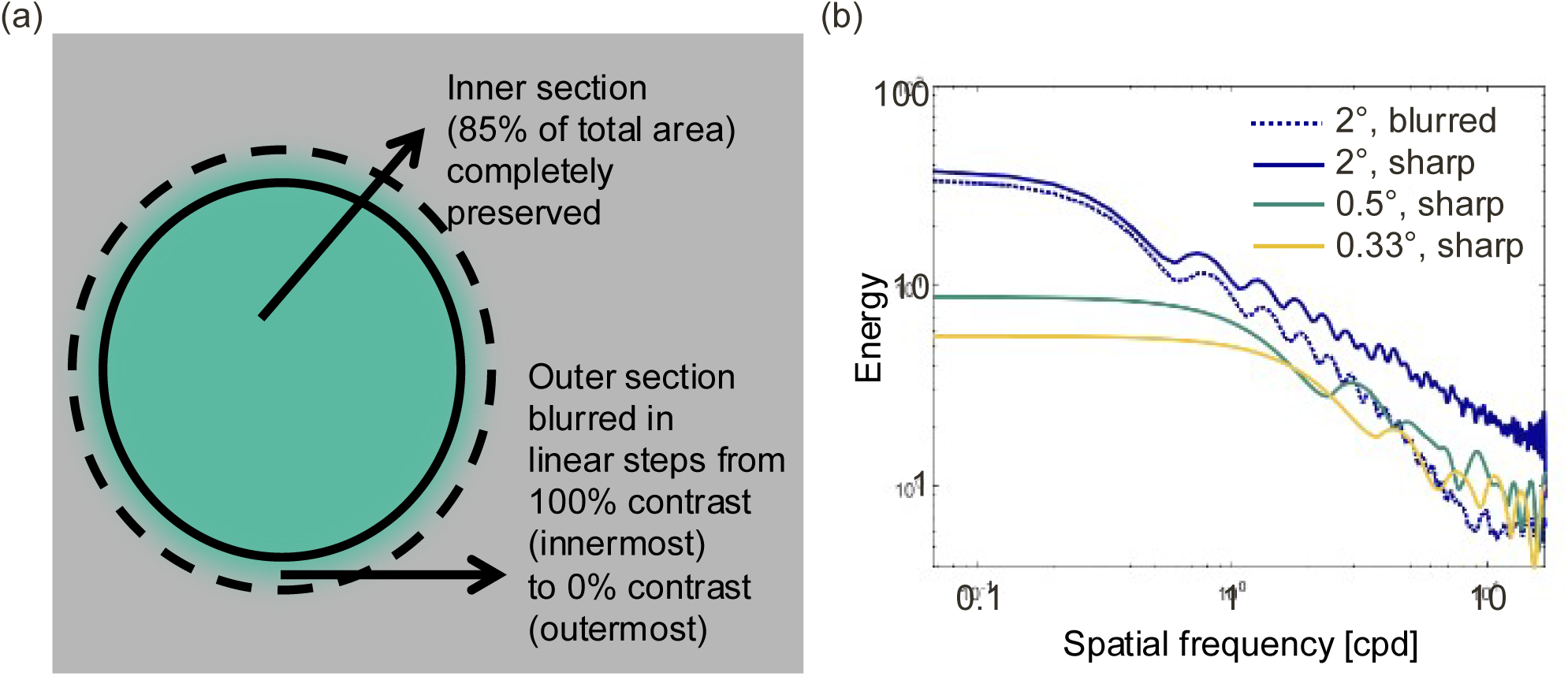
Schematic representation of stimulus edge blurring (a) and stimulus energy as a function of spatial frequency (b) for blurred and sharp edge stimuli of different sizes. The contrast of the central region (labelled “inner section” in panel A) is set to identical contrast levels for all discs depicted in panel B. Note that the higher frequency content of a 2° disc stimulus depends overwhelmingly on its edge – once the edge is blurred by adding a linear contrast ramp over its outer area, the disc maintains a highly similar amount of energy in the lower spatial frequencies (< 0.2 cycles per degree) but its higher spatial frequency content (>2 cycles per degree) has dropped off significantly and is now within the range of the smaller 0.5° and 0.33° discs.

### Statistical Data Analysis

We used linear mixed effect models on log-transformed contrast adjustment data and therefore report effect sizes as odds ratios (ORs). Analyses were implemented in R 4.3.1 (R Core Team, 2024) using the following packages: lmerTest 3.1-3 (Kuznetsova et al., 2017) emmeans 1.10 (Lenth et al., 2025), lmeresampler 0.2.4 (Loy et al., 2023), DHARMa 0.4.6 (Hartig et al., 2024) and simr (Green & MacLeod, 2016). e report further details on statistical analyses in Supplementary Materials where we also present statistical sensitivity analyses for our models, based on Kumle et al.’s (2021) recommendations of evaluating sensitivity for LMEMs by reducing fixed effects by 15% and simulating the analyses for different sample sizes re-drawn from the original sample. Supplementary Materials also contain the full properties of best-fitting models and pairwise post-hoc tests for our effects.

To further confirm whether the increase in adjustment with decreasing stimulus size differed between chromatic and achromatic stimuli, we conducted a Bayesian model comparison using the anovaBF function, part of the BayesFactor (Morey et al., 2024) package in R. The Bayesian factor estimates for each model were based on 10000 iterations. Our interpretations of Bayes factors are based on Kass & Raftery (1995) and we refer to values BF_10_ < 1 as evidence against hypothesis, 1-3 as weak, 3-20 as moderate and > 20 as strong evidence for the hypothesis.

### Transparency and Openness

We have reported how we determined out sample size, and we also report all data exclusions, all manipulations and all measures in the study. Data collection took part between December 2021 and March 2022. All data and code required to replicate the experiment and analysis have been made available on OSF (osf.io/jmght). This study was not preregistered.

## Results

### 1. Comparing Desaturation with Size Between Chromatic and Luminance Stimuli

We analyzed the chromatic and luminance data using linear mixed effect models. For color, we could only achieve a threefold increase in standard contrast, whilst for luminance we also tested standards that were nine times higher, so we fitted two separate models to these datasets.

First, a linear mixed effects model was fitted to chromatic and luminance data, predicting adjustment as a function of size, color and contrast level (baseline level and 3x higher contrast). The best fitting model with random by-participant intercepts explained r^2^=0.589 variance, with fixed effects accounting for *r ^2^*= .432. Figure 4a shows the prominent reduction of perceived contrast for smaller stimuli (0.5° vs 2°: OR 1.132, 95% CI 1.107-1.157, *t*(491) = 9.740, *p* <. 001; 0.33° vs. 0.5°: OR 1.081, 95% CI 1.054-1.108, *t*(491) = 5.779, *p* < .001). A significant interaction between size and contrast level was present, with desaturation significantly reduced for higher contrast stimuli at 0.33° (OR 1.073, 95% CI 1.031-1.115, *t*(491) = 3.596, *p* =.003). Desaturation with size also significantly differed across colors, with a higher loss of perceived contrast for positive luminance polarity at 0.5° against all the hues except blue – a loss which at the smallest, 0.33° size had become more uniform across colors (see Supplementary Materials for all the post-hoc tests).

**Figure 4.**
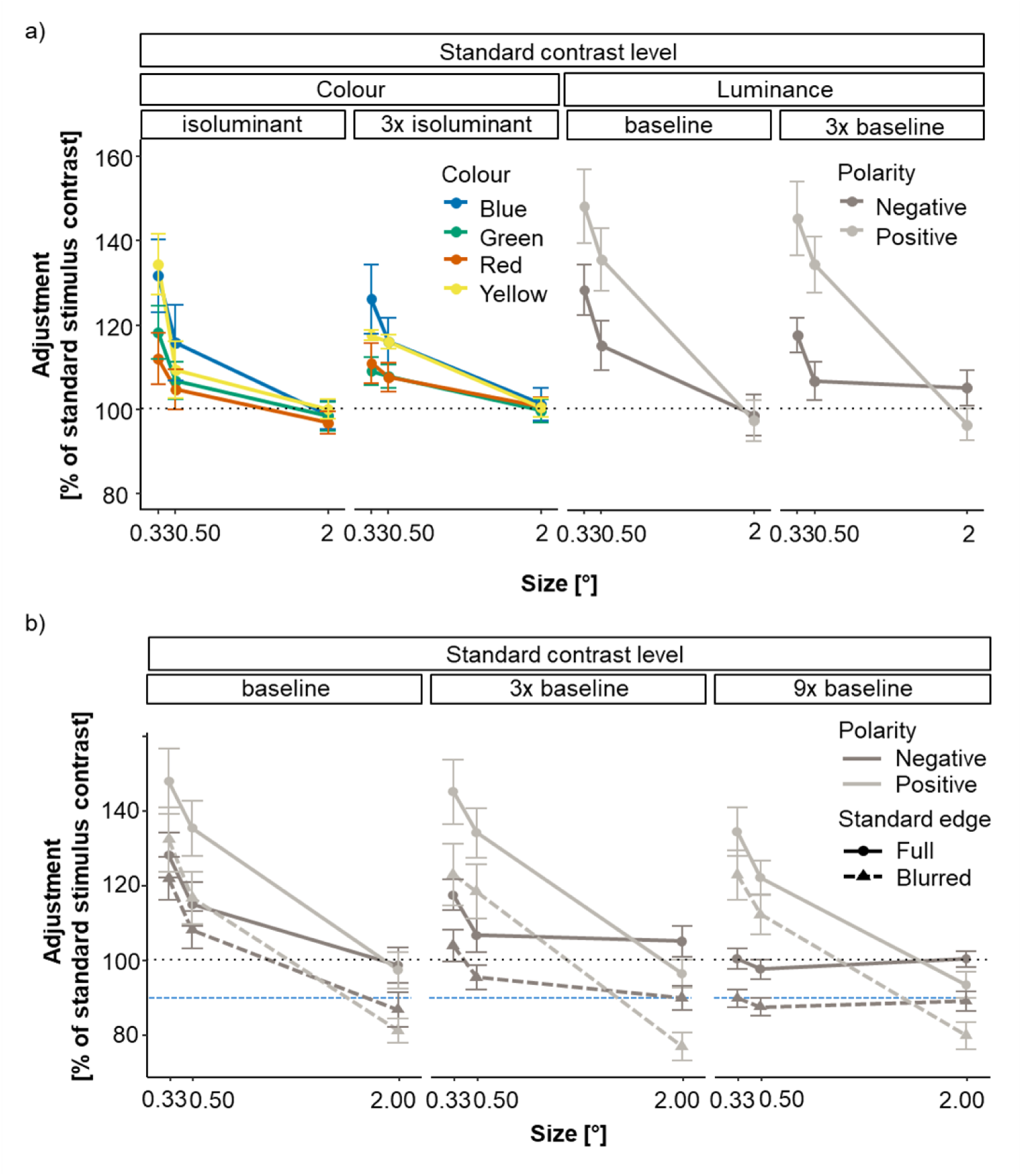
Perceived contrast for chromatic and luminance-defined stimuli of different sizes. (a) Isoluminant color stimuli and luminance-defined stimuli matched to suprathreshold standards of equal salience (i.e., at a relatively well-saturated level within the monitor gamut) as well as to standards 3 times higher in contrast. (b) Luminance-defined stimuli as a function of stimulus size, depicting matches for a suprathreshold contrast level (left) and for three (middle) and nine (right) times higher contrast for sharp (full line) and blurred edge (dashed line) stimuli. Note that the left and middle panels for sharp edge stimuli are identical to the luminance-defined conditions in (a) and are replotted to facilitate comparisons. To compare the extent of desaturation between chromatic and luminance-defined stimuli, we represent saturation in units of standard stimulus contrast. Therefore, an adjustment value of 200% would indicate twice the amount of contrast in the standard stimulus was required for the comparison stimulus to achieve the same perceived saturation. The dotted line at 100% indicates the contrast of the central region of the standard stimulus, which in the case of sharp-edge standards is identical to their full contrast. Dashed blue line indicates absolute contrast value for blurred edge stimuli (92.5%). Error bars denote 2 standard errors of the between-subject mean.

A linear mixed effects model was also fitted to the achromatic data (Figure 4b), predicting contrast adjustment as a function of size, contrast level, luminance polarity and edge definition. The best-fitting model explained *r*^2^ = 0.633 through the fixed effects, rising to *r*^2^ = 0.745 with the inclusion of random by-participant intercepts. The three-way interaction between size, contrast level and polarity was significant -as the standard contrast was tripled, desaturation of 0.33° discs was lowered for negative luminance polarity only (3x contrast vs. baseline: OR 1.124, 95% CI 1.048-1.200, *t*(369) = 3.403, *p* = .028). With a further tripling of luminance contrast, perceived contrast was even less affected by stimulus size for negative luminance polarity discs (9x vs. 3x contrast: OR 1.162, 95% CI 1.084-1.240, *t*(369) = 4.353, *p* < .001). In fact, at 9 times higher contrast a flat profile is observed for negative polarity stimuli, with a lack of statistically robust differences (2° vs 0.5°: OR 1.024, 95% CI 0.955-1.093, *t*(369) = 0.676, *p* = 1.00; 0.5° vs. 0.33°: OR 0.972, 95% CI 0.906-1.038, *t*(369) = -0.814, *p*=1.00; see Supplementary Materials for further details of the post-hoc tests).

As can be seen from Figure 4b, contrast adjustments were consistently higher when luminance-defined stimuli were matched to a sharp edge standard than to a standard with a blurred edge. The effect of blurring surpasses the difference in overall contrast between the standard sharp-edged and blurred-edge stimulus (*t*-test against contrast of 92.5% *t*(9) = -10.848, *p* < .001, *M* = 83.778% 95% CI 81.354’% 86.201%, *d* = -3.43), demonstrating the importance of edges in driving our perceptual experience of contrast (see also Figure 5 for convergent findings from our 2^nd^ experiment).

**Figure 5.**
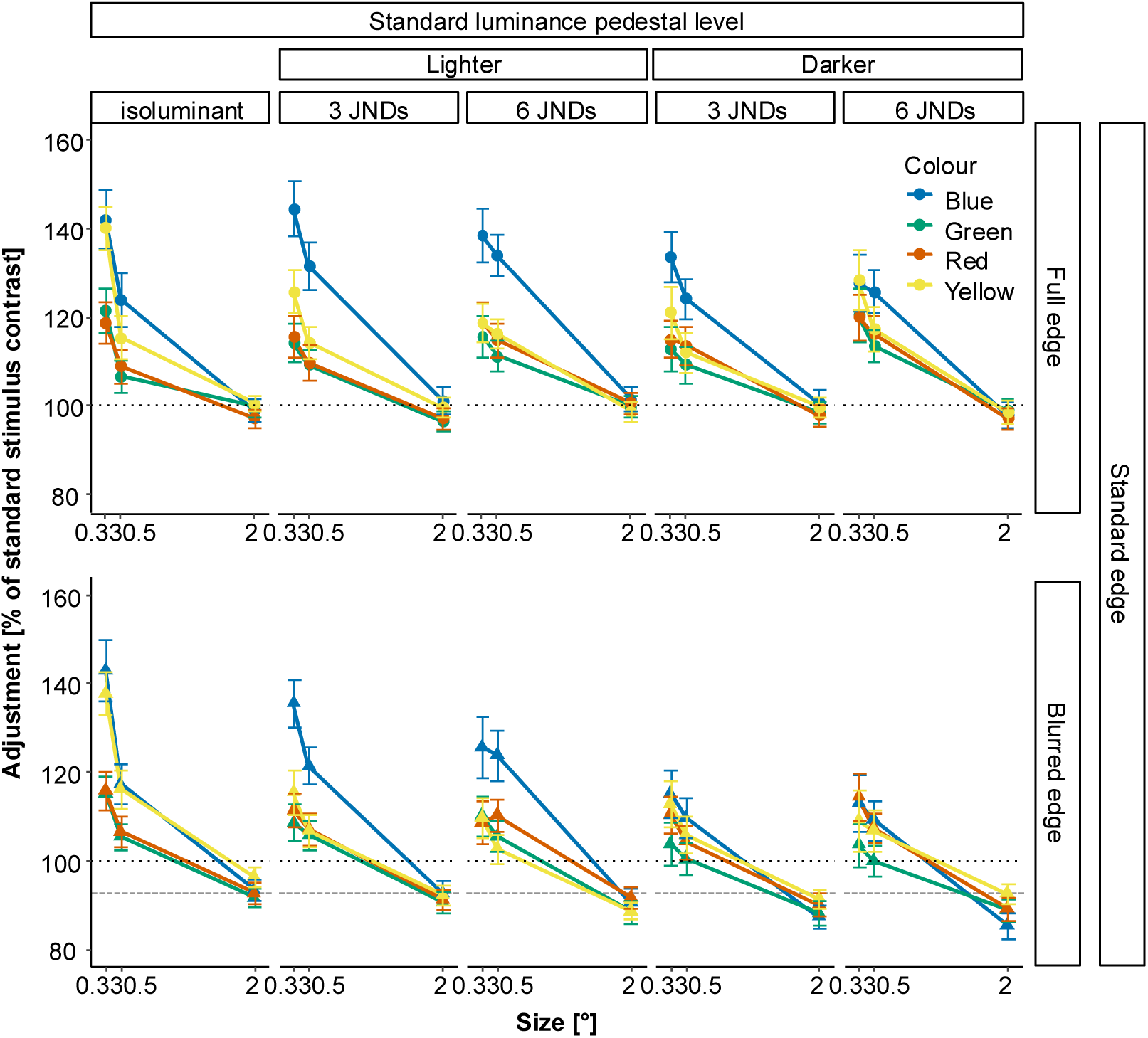
Perceived contrast for colored discs of different size, presented with and without luminance pedestal, matched to either a sharp-edged or blurred-edged standard stimulus. Perceived contrast is presented with and without luminance contrast (i.e.., lighter: 3 JND+ and 6 JND+ and Darker: 3 JND- and 6JND-, with + standing for positive and – for negative luminance polarity), adjustment is expressed as percentage of standard stimulus contrast, as in Figure 4. The dotted line at 100% indicates the contrast of the central region of the standard stimulus, which in the case of sharp-edge standards is identical to their full contrast. In the lower panel, an additional grey line indicates the actual contrast of the blurred edge stimulus (92.5%). Error bars represent 2 standard errors of the between-subject mean.

As outlined in the methods section, to confirm whether the increase in adjustment with decreasing stimulus size differed between chromatic and achromatic stimuli, we conducted a Bayesian model comparison. Firstly, as expected, we found evidence against a model predicting adjustment performance based on the colour or luminance mechanism alone (BF_10_ = 0.12). Furthermore, whilst a model based on size alone provided extremely strong evidence for desaturation (BF_10_ = 6.22 × 10⁴³), this was slightly reduced with the addition of the mechanism factor (BF_10_ = 9.06 × 10⁴²), thus suggesting the desaturation with size is comparable between mechanisms. Addition of the interaction between the factors of size and mechanism increased the evidence compared to a non-interactive model, but this increase was weak compared to the size only model (BF = 1.36). Taken together, the evidence suggests that the effect of desaturation with a decrease in size is largely comparable between color and luminance mechanisms.

### 2. Desaturation with a Reduction of Stimulus Size for Light and Dark Colored Discs with Blurred or Sharp Edge

A linear mixed-effects model was fitted to the data from the 2^nd^ experiment, predicting perceived contrast as a function of size, color, luminance contrast pedestal and edge definition (for full details, see Supplementary Materials). There was a series of two-way interactions between the factors, with the best-fitting model explaining about half of the variance in the dataset (Marginal *R*^2^ = 0.537, Conditional *R*^2^ = 0.663). First, an interaction between color and size replicated our previous findings: as shown in Figure 5, reduction of stimulus size led to desaturation: the amount of contrast required to achieve equal saturation of the comparison stimulus increased as stimulus size decreased (2° vs. 0.5°: OR 0.848, 95% CI 0.840-0.856, *t*(2297) = -34.688, *p* < .001; 0.5° vs. 0.33°: OR 0.944, 95% CI 0.935-0.953, *t*(2297) = -11.854, *p* < .001). Desaturation was highest for blue (vs. yellow: OR 1.053, 95% CI 1.041-1.065, *t*(2297) = 9.145, *p* < .001), followed by yellow (vs green: OR 1.048, 95% CI 1.037-1.059, *t*(2297) = 8.806, *p* < .001; for further post-hoc tests, see Supplementary Materials).

Luminance pedestals interacted with color, as well as with size. Addition of luminance pedestals had a very small effect on desaturation of red or green. However, yellow and blue were both significantly and systematically affected. For yellow, all of the luminance pedestals led to less desaturation compared to isoluminance, without significant differences between the different pedestals. For blue, desaturation was reduced only by negative luminance pedestals (see Supplementary Materials for all post-hoc tests of this interaction). Interaction of luminance pedestal with size was such that pedestals led to no significant changes in perceived contrast for the 2° stimulus, but produced significantly less desaturation compared to isoluminance for the 0.33° stimulus (again, see Suppl. Materials for all the details of the post-hoc tests).

Together with our predictions of less desaturation with the addition of luminance pedestals, we also predicted blurring of the standard stimulus edge to decrease perceived contrast. This would mean that less contrast would be required to find a match with a blurred-edge standard, compared to a sharp-edge standard. There was also a significant interaction between edge definition and luminance pedestal: for blurred-edge discs, the isoluminant condition required more contrast to achieve equal salience in relation to all conditions with luminance pedestals except for the positive pedestal of 3 JNDs, which similarly to isoluminance required significantly more contrast than the negative pedestal conditions (again, see Suppl. Materials for all the details of the post-hoc tests). This is in line with our prediction that blurring the edge would particularly affect perceived contrast once color is combined with luminance information.

## Discussion

We demonstrate that parafoveal desaturation with a reduction in stimulus size is characteristic of general contrast processing by the visual system, rather than specific to color. This may appear counterintuitive, as it is generally accepted that color perception is a specialization of foveal vision, despite extensive evidence that this is not the case (e.g. Tyler, 2015). Here, we show that once achromatic stimuli are perceptually matched in contrast with chromatic stimuli, they exhibit the same changes in appearance with a reduction in stimulus size as chromatic stimuli. We demonstrate that this effect can be reduced when overall contrast of chromatic stimuli is sufficiently increased or shifted towards higher spatial frequency channels through the addition of luminance pedestals. In fact, at contrast levels tested in this study, reduction in perceived contrast with a decrease in stimulus size is eliminated for negative but not for positive luminance polarity stimuli. It is not possible to achieve sufficient contrast levels within a monitor’s gamut to alleviate the desaturation with size for purely chromatic stimuli. We also show that the extent of desaturation differs between the two chromatic mechanisms and that it can be reduced for the most impacted blue-yellow mechanism when presented in combination with luminance contrast pedestals. These findings highlight that general contrast mechanisms can explain loss of perceived contrast as experienced in the parafovea with a reduction in stimulus size.

Contrast constancy theory (Kulikowski, 1976) account for our findings fairly well. For positive luminance contrasts detection threshold is higher (Lu & Sperling, 2012), which would make low contrast stimuli closer to threshold and thus reduced in perceived contrast, especially when parafoveally presented. Contrast constancy can be achieved when physical contrast is sufficiently higher than threshold. For low to medium contrast gratings or Gabors, the point of constancy depends on spatial frequency (Georgeson & Sullivan, 1975) but for disc stimuli it depends on the length of an edge, with the decrease in sensitivity as size reduces well explained by probability summation over multiple edge detectors, following the envelope of the CSF (Ashraf et al., 2023). When contrast is at threshold, only the most sensitive channel processes the information (Kulikowski, 2003). When contrast is sufficiently increased, it breaches the contrast sensitivity limitations of other channels and they begin to contribute to contrast perception. Our investigation of perceived contrast with blurred edge standards (i.e., reduced high-frequency information) confirms this: matches of full-edge comparison discs to blurred edge standards are affected across all sizes, with the effect on appearance far exceeding the overall change in standard stimulus contrast (see Fig. 4).

Similar findings on perceived contrast across chromatic and achromatic stimuli in the periphery are reported by Jiang and colleagues (2022). Their observers compared perceived contrast of a peripheral stimulus to the foveally presented standard at varying suprathreshold contrast levels until the point of objective equality (PSE) was achieved. Whilst constancy was achieved at higher contrasts, the upper range of spatial frequencies appeared less contrasty than predicted by the contrast constancy model when the standard was presented at low or middle contrast (for consistent findings of deviations from contrast constancy, see Ashraf et al., 2022). This suggests a ‘hybrid model’ of contrast constancy, where constancy is not entirely achievable for higher-frequency stimuli at lower or mid-ranges of contrast. Such a hybrid model is in line with our own observations here. Put together, these findings point towards desaturation as a consequence of low-level contrast bottlenecks. Our smaller stimuli appear desaturated as the general contrast sensitivity for high spatial frequencies is low for chromatic information (Mullen, 1985) and decreases further with eccentricity (Anderson et al., 1991). Due to their small size (0.33° and smaller), their energy distribution cannot be effectively processed in the parafovea. Saturation for larger discs depends on summed information from a larger number of channels and is therefore preserved at these eccentricities.

Luminance contrast pedestals play an important role in preserving saturation for blue and yellow stimuli. For yellow, change is observed even with lower positive and negative luminance pedestal magnitudes, whereas for blue, only negative polarity pedestals increase perceived contrast and thus reduce desaturation with size. The blue-yellow mechanism CSF has a lower spatial frequency cut off than the red-green mechanism (Wuerger et al., 2020). The actual number of spatial channels within each of the mechanisms has not yet been definitely established (Peterzell & Teller, 2000), but previous research suggests three channels for red-green (Losada & Mullen, 1994) and two channels for blue-yellow, matching the sensitivity of the two lowest spatial frequency luminance channels (Humanski & Wilson, 1992). Inclusion of additional luminance contrast energy exciting the range of higher spatial frequency channels within the luminance CSF envelope would be expected to decrease detection threshold and hence increase perceived contrast. Different impact of luminance contrast on yellow compared to blue may be due to differently spatially tuned Short-wavelength-ON (responding to blue) and Short-wavelength-OFF mechanisms (responding to yellow; Hardman & Martinovic, 2021).

Finally, we need to consider if our results are influenced by eye movements. This is unlikely for several reasons. Our experienced psychophysical observers were instructed to only perform the task when steadily fixating, and produced patterns of desaturation consistent with low-level influences on perception, as described above. Jiang et al. (2022) obtained highly convergent patterns of results with a very similar adjustment task. In fact, if participants were foveating the stimuli consecutively, they would have produced flat lines in both our studies, without any effects of eccentricity.

In conclusion, perceptual experience of color saturation is comparable to the well-explored contrast perception of luminance stimuli. This means that further insights into low-level mechanisms of contrast processing, across both luminance and color, can better inform our understanding of high-level color properties such as saturation and how these can be accurately represented in models of color appearance. Similar research programs have already led to a collection of large-scale threshold and suprathreshold datasets whose modelling not only has practical applications in the domain of display technologies (e.g., Ashraf et al., 2024) but can also lead to interesting predictions in relation to human perception mechanisms. For example, they would accurately predict our current findings, dispelling the often-accepted myth that it is only color whose appearance is affected when presented in the periphery.

## Notes

### Competing Interest Statement

The authors have declared no competing interest.

### Summary of Updates

Updates made to the introduction and new analyses (bayesian statistics) added.

