## Supplementary Materials for "Color Desaturation in the Periphery is Explained by General Mechanisms of Contrast Sensitivity and Constancy"

### SUPPLEMENTRY MATERIALS 1 – STATISTICAL ANALYSES

#### **1.1 Experiment 1**: Comparing desaturation with size between chromatic and luminance stimuli

#### 1.1.1. CHROMATIC AND ACHROMATIC DATA (baseline and 3x contrast)

**Best fitting linear mixed effect model to chromatic and achromatic data combined for contrast expressed as % of standard stimulus**, with fixed effects of size (2° - reference level, 0.5° and 0.33°), contrast level (at baseline setting and 3 times higher), colour (red – reference level, green, blue, yellow, positive luminance polarity and negative luminance polarity) and edge (sharp – reference level, or blurred) and random by-participant intercepts only.

Model: normalised contrast (in %) ~ size+contrast level+colour+ size:contrast level + size:colour + 1|participant

| **Fixed Effects** | **Estimate** | **t value** | **p value** | **95% CI** |
|---|---|---|---|---|
| Intercept | 110 | 193.528 | **<.001*** ** | 106-115 |
| 0.5 vs. 2 | 15.7 | 9.954 | **<.001*** ** | 10.5-20.8 |
| 0.33 vs. 2 | 25.8 | 15.861 | **<.001*** ** | 18.9-32.6 |
| 3x vs. baseline contrast | -2.97 | -1.772 | .077 | -7.36- 1.51 |
| Green vs. red | 1.25 | 0.606 | 0.545 | -1.25 – 3.78 |
| Blue vs. Red | 9.51 | 3.366 | **<.001*** ** | 4.01-15.3 |
| Yellow vs. Red | 7.60 | 2.900 | **0.004** ** | 3.59-11.9 |
| Positive polarity vs. Red | 11.9 | 4.173 | **<.001*** ** | 5.25-18.5 |
| Negative polarity vs. Red | 1.16 | 0.245 | 0.807 | -1.00 – 3.40 |
| 0.5 vs. 2 / 3x vs. bas. contrast | -2.81 | -0.756 | 0.450 | -8.21 – 2.39 |
| 0.33 vs. 2 / 3x vs bas. contrast | -10.8 | -3.191 | **0.0015** ** | -15.5- -6.22 |
| 0.5 vs. 2 / Green vs. Red | 0.809 | 0.174 | 0.862 | -4.95-6.78 |

| | | | | |
|---|---|---|---|---|
| 0.33 vs. 2 / Green vs. Red | 1.88 | 0.366 | 0.715 | -4.77-8.69 |
| 0.5 vs. 2 / Blue vs. Red | 8.59 | 1.131 | 0.259 | -1.57-19.3 |
| 0.33 vs. 2 / Blue vs. Red | 16.3 | 2.360 | **0.019*** | 4.14-28.9 |
| 0.5 vs. 2 / Yellow vs. Red | 5.01 | 0.674 | 0.500 | -2.79 – 13.3 |
| 0.33 vs. 2 / Yellow vs. Red | 13.0 | 1.983 | **0.048*** | 3.52-23.2 |
| 0.5 vs. 2 / Positive P. vs. Red | 30.9 | 6.123 | **<.001***** | 22.5-39.6 |
| 0.33 vs. 2 / Positive P. vs Red | 36.6 | 6.830 | **<.001***** | 24.2-49.3 |
| 0.5 vs. 2 / Negative P. vs. Red | 3.83 | 0.818 | 0.414 | -0.713 – 8.60 |
| 0.33 vs. 2 / Negative P. vs Red | 10.1 | 2.085 | **0.038*** | 3.66-16.9 |
| *Random effects* | | | | |
| Residual variance, $\sigma^2$ = 194.71 | | | | |
| Participant variance 75.43 | | | | |
| ICC 0.28 | | | | |
| N participant 10, 480 Observations in total | | | | |
| Marginal $R^2$ / Conditional $R^2$ 0.432 / 0.589 | | | | |

To evaluate the distribution of residuals we used simulation-based residual diagnostics implemented in the DHARMa R package. The residuals for the initial model fit to the full dataset averaged across trials suffered from underdispersion, due to the symmetric contrast matches for 2° stimuli being much less dispersed than the other stimulus sizes. The main issue of underdispersion is the loss of power, inflating type II error and making the statistical tests more conservative. To correct the distribution (which for a zero-bound variable such as contrast matching will be right-skewed), we fitted the model to log-transformed data and report the statistical results (F and t-values and p-values) from this model, while the estimates and 95% CIs are reported from the bootstrapped model (with 1000 resamples of participants, as implemented in R package lmeresampler). Corresponding to the estimates, we report residual and participant variance from the non-log corrected model.

As evaluated by backward-reduced model fitting implemented in function *step* from R package lmerTest, the interactions that could be removed from the model were as follows: 3-way interaction ($F(10,470)=0.985, p=0.456$) and the 2-way interactions between contrast level and colour ($F(5,470)=0.886, p=0.490$). The following 2-way interactions could not be removed from the model: size by contrast level (F(2,470)=6.486,p=.002) and size by colour (F(10.470)=8.914,p<.001).

### POST-HOC ANALYSES FOR 2-WAY INTERACTIONS

Post-hoc analyses were performed using *emmeans* package for R. We used the multivariate t distribution adjustment ('mvt') correction method for multiple comparisons for a series of t-tests across all the levels of the interaction.

#### *Interaction Between Size and Contrast Level*

The differences between contrast levels only emerged for the 0.33 stimulus, for which desaturation was more prominent for stimuli at the lower level of contrast:

2 deg: Baseline vs. 3x higher contrast – OR 0.984, 95%CI 0.946-1.022, t(491) = -0.82, p = .948

0.5 deg: Baseline vs. 3x higher contrast – OR 1.004, 95%CI 0.965-1.043,t(491)=0.277, p=1.00

0.33 deg: Baseline vs. 3x higher contrast – OR 1.073, 95%CI 1.031-1.115,t(491)=3.596, **p=.003****

#### *Interaction Between Colour and Size*

For red, we do not note a statistically significant desaturation with size:

Red: 2 vs. 0.5, OR 0.933, 95%CI 0.860-1.006, t(491) = -1.787, p= 0.8935

Red: 2 vs. 0.33, OR 0.893, 95%CI 0.825-0.961, t(491) = -2.917 p= 0.1520

Red: 0.5 vs. 0.33, OR 0.957, 95%CI 0.884-1.030, t(491) = -1.130 p= 0.9990

For all other colours, there is a statistically significant desaturation for at least the smallest stimulus:

Green: 2 vs. 0.5, OR 0.924, 95%CI 0.853-0.995, t(491) = -2.027, p= 0.753

Green: 2 vs. 0.33, OR 0.875, 95%CI 0.808-0.942, t(491) = -3.423, **p= 0.0334***

Green: 0.5 vs. 0.33, OR 0.947, 95%CI 0.875-1.019, t(491) = -1.396, p= 0.989

Blue: 2 vs. 0.5, OR 0.878, 95%CI 0.811-0.945, t(491) = -3.352, **p= 0.0414***

Blue: 2 vs. 0.33, OR 0.786, 95%CI 0.726 – 0.846, t(491) = -6.183, **p<.001*****

Blue: 0.5 vs. 0.33, OR 0.896, 95%CI 0.828-0.964, t(491) = -2.831, p= 0.189

Yellow: 2 vs. 0.5, OR 0.899, 95%CI 0.830-0.968, t(491) = -2.721, p= 0.245

Yellow: 2 vs. 0.33, OR 0.802, 95%CI 0.741-0.863, t(491) = -5.662, **p <.001*****

Yellow: 0.5 vs. 0.33, OR 0.892, 95%CI 0.824-0.960, t(491) = -2.941, p= 0.142

Positive Polarity: 2 vs. 0.5, OR 0.701, 95%CI 0.663-0.739, t(491) = -12.905, **p<.001*****

Positive Polarity: 2 vs. 0.33, OR 0.649, 95%CI 0.614-0.684, t(491) = -15.701, **p<.001*****

Positive Polarity: 0.5 vs. 0.33, OR 0.926, 95%CI 0.876-0.976, t(491) = -2.796, p= 0.205

Negative polarity: 2 vs. 0.5, OR 0.898, 95%CI 0.850-0.946, t(491) = -3.913, **p= 0.006****

Negative polarity: 2 vs. 0.33, OR 0.810, 95%CI 0.766-0.854, t(491) = -7.659,  **p <.001*****

Negative polarity: 0.5 vs. 0.33, OR 0.902, 95%CI 0.853-0.951, t(491) =  -3.745,  **p= 0.0105***

**SENSITIVITY ANALYSES FOR THE OBSERVED EFFECTS**

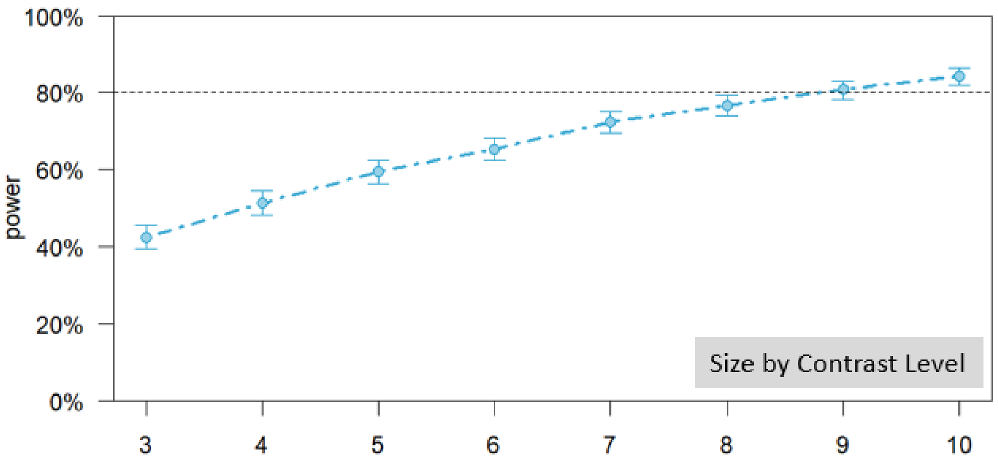

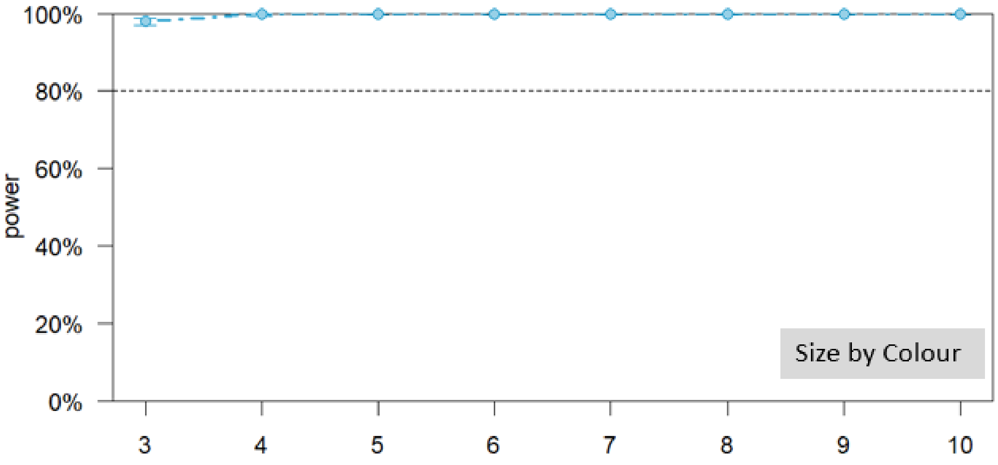

Sensitivity analyses were performed by resampling the number of participants (1000 simulations) and evaluating the observed power for obtaining the statistically significant fixed effects in the best fitting model, using the R package *simr*. Sufficient power for those effects can be observed with 8 out of 10 participants for the interaction of size with contrast level, and only 3 participants are needed to reach 80% power (indicated by a dotted line) for the highly significant interaction between size and colour. We also evaluated the power to detect the observed effects if they were reduced by 15% (Kumle et al., 2021), which was again very high (100%; CI 99.63-100%).

**1.1.2 ACHROMATIC DATA (baseline, 3x and 9x contrast)**

**Best fitting linear mixed effect model to achromatic data for contrast expressed as % of standard stimulus**, with fixed effects of size (2° - reference level, 0.5° and 0.33°), contrast level (at baseline – reference level, 3 times higher and 9 times higher), luminance polarity (positive – reference level, and negative) and edge definition (sharp – reference level, or blurred) and random intercepts only.

Model: normalised contrast (in %) ~ size+contrast level+polarity+ edge definition + size:contrast level + size: polarity + contrast level: polarity + size:contrast level:polarity + 1|participant

| **Fixed Effects** | **Estimate** | **t value** | **p value** | **95% CI** |
|---|---|---|---|---|
| Intercept | 109 | 202.829 | **<.001*** ** | 104-113 |
| 0.5 vs. 2 | 21.3 | 14.673 | **<.001*** ** | 15.5-27.2 |
| 0.33 vs. 2 | 31.2 | 20.483 | **<.001*** ** | 22.9-39.7 |
| 3x vs. baseline contrast | -4.80 | -2.889 | 0.004 | -9.78 – 0.038 |
| 9x vs. baseline contrast | -11.7 | -7.254 | **<.001*** ** | -20.0- -3.62 |
| Positive vs. Negative polarity | -12.7 | -8.792 | **<.001*** ** | -18.7 - -6.91 |
| Blurred vs. sharp edge | -13.4 | -11.355 | **<.001*** ** | -15.3 - -11.4 |
| 0.5 vs. 2 / 3x vs. bas. contrast | -6.28 | -1.686 | 0.093 | -14.6 – 1.86 |
| 0.33 vs. 2 / 3x vs bas. contrast | -11.4 | -2.744 | **0.006** ** | -18.2 - -4.69 |
| 0.5 vs. 2 / 9x vs. bas. contrast | -13.7 | -3.595 | **<.001*** ** | -20.2 - -7.25 |
| 0.33 vs. 2 / 9x vs bas. contrast | -20.6 | -5.069 | **<.001*** ** | -26.6 - -14.6 |
| 0.5 vs. 2 / Pos. vs. Negative | -29.0 | -9.991 | **<.001*** ** | -37.4 - -20.8 |
| 0.33 vs. 2 / Pos. vs. Negative | -31.6 | -10.094 | **<.001*** ** | -43.4 - -20.1 |
| 3x vs. base / Pos. vs. Negative | -3.85 | -0.946 | 0.345 | -10.1 – 2.32 |
| 9x vs. base / Pos. vs. Negative | -8.02 | -3.038 | **0.002** ** | -15.1 - -0.78 |
| 0.5 vs. 2 / 3x vs. bas. Contrast / Positive vs. Negative | -18.2 | -2.581 | **0.010*** | -24.9 - -11.4 |
| 0.33 vs. 2/3x vs. bas. Contrast / Positive vs. Negative | -15.7 | -2.283 | **0.023*** | -24.7 - -6.95 |
| 0.5 vs. 2 / 9x vs. bas. Contrast / Positive vs. Negative | -14.9 | -2.448 | **0.015*** | -25.6 - -4.33 |

| | | | | |
|---|---|---|---|---|
| 0.33 vs. 2/9x vs. bas. Contrast / Positive vs. Negative | -23.3 | -3.595 | **<.001*** ** | -38.1 - -8.39 |
| *Random effects* | | | | |
| Residual variance, $\sigma^2$ = 161.29 | | | | |
| Participant variance 76.05 | | | | |
| ICC 0.31 | | | | |
| N participant 10, 360 Observations in total | | | | |
| Marginal $R^2$ / Conditional $R^2$ 0.633 / 0.745 | | | | |

To evaluate the distribution of residuals we used simulation-based residual diagnostics implemented in the DHARMa R package. Again, the residuals for the model fit to the full dataset averaged across trials suffered from underdispersion, due to the contrast matches for 2° stimuli and stimuli at 9x contrast being much less dispersed than the other conditions (i.e., being very closely aligned to standard stimulus contrast). The main issue of underdispersion is the loss of power, inflating type II error and making the statistical tests more conservative. To correct the distribution (which for a zero-bound variable such as contrast matching will be right-skewed), we fitted the model to log-transformed data and report the statistical results (F and t-values and p-values) from this model, while the estimates and 95% CIs are reported from the bootstrapped model (with 1000 resamples of participants, as implemented in R package lmeresampler). Corresponding to the estimates, we report residual and participant variance from the non-log corrected model.

As evaluated by backward-reduced model fitting implemented in function *step* from R package lmerTest, The interactions that could be removed from the model were as follows: 4-way interaction ($F(4,350)=0.130, p=0.971$), 3-way interactions between size, contrast level and edge definition ($F(4,350)=0.157, p=.960$), contrast level, polarity and edge definition ($F(2,350)=0.686, p=.504$) and size, polarity and edge definition ($F(2,350)=0.0904, p=0.914$) and the 2-way interactions between size and edge definition ($F(2,350)=0.306, p=0.737$), contrast level and edge definition ($F(2,350) = 1.303$, $p=.273$) and polarity and edge definition ($F(1,350)=3.723$, $p=.0545$). The following effects could not be removed from the model: 3-way interaction between size, contrast level, and polarity ($F(4,350)=2.7595, p=.0277$) and edge ($F(1,350)=99.761, p<.001$).

### POST-HOC ANALYSES FOR THE 3-WAY INTERACTION BETWEEN SIZE, CONTRAST LEVEL AND LUMINANCE POLARITY

Post-hoc analyses were performed using *emmeans* package for R. We used the multivariate t distribution adjustment ('mvt') correction method for multiple comparisons for a series of t-tests across all the levels of the interaction.

For the positive luminance polarity, the loss of saturation with size appeared to be equal across different contrast levels (all ts < 2.262, all ps > 0.512).

On the contrary, for the negative luminance polarity there was significantly less desaturation between 9x higher contrast and contrast at baseline level, both at 0.5° (OR 1.196, 95%CI 1.115-1.277, t(369) = 5.203, **p<.001****) and 0.33° (OR 1.306, 95%CI 1.218-1.394, t(369) = 7.756, **p<.001****). There was also a significant decrease in desaturation between 3x higher contrast and contrast at baseline for the 0.33° stimulus (OR 1.124, 95%CI 1.048-1.200, t(369) = 3.403, **p = 0.028***). Finally, the desaturation profile was significantly flatter at 0.33° for 9x higher contrast compared to 3x higher contrast too (OR 1.162, 95%CI 1.084-1.240, t(369) = 4.353, **p<.001*****). All other tests were insignificant (ts < 2.664, ps>.235).

### SENSITIVITY ANALYSES FOR THE OBSERVED EFFECTS

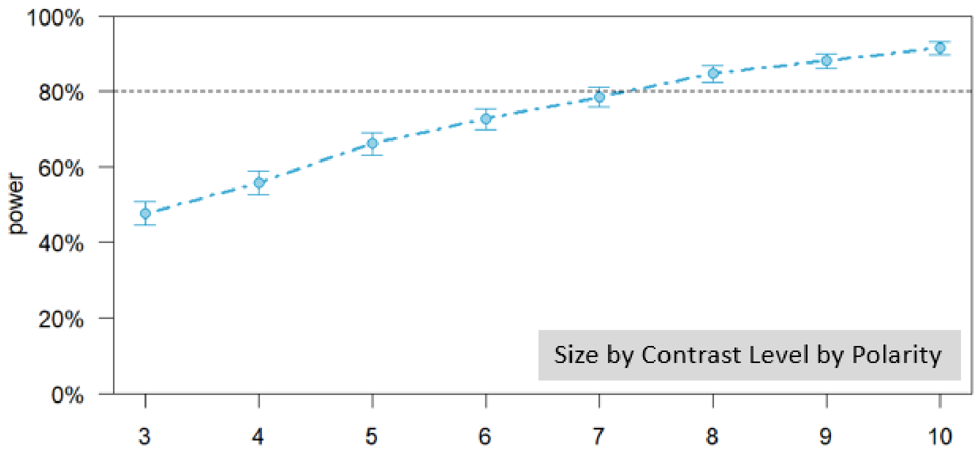

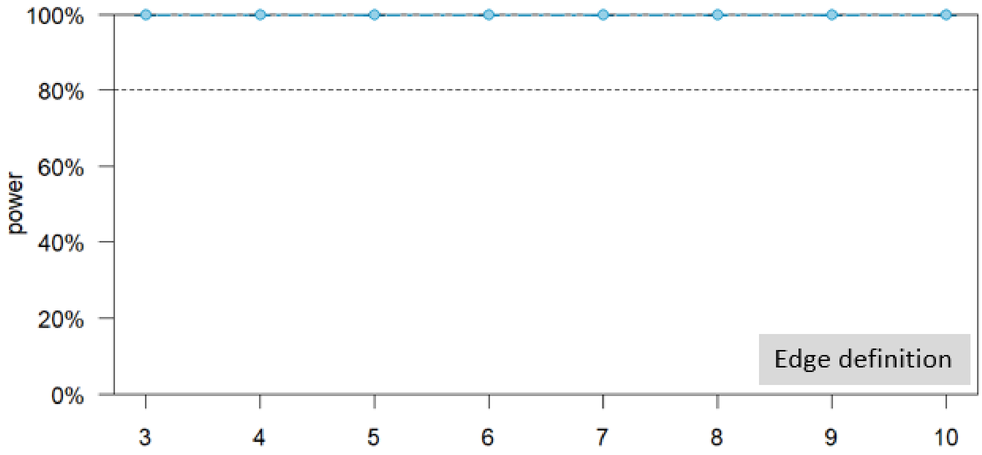

Sensitivity analyses were performed by resampling the number of participants (1000 simulations) and evaluating the observed power for obtaining the statistically significant fixed effects in the best fitting model, using the R package *simr*. Power exceeds 80% with 8 out of 10 participants for the 3-way interaction of size, contrast level and luminance polarity, while 3 participants are sufficient (indicated by a dotted line) for 100% power for the very strong and robust fixed effect of edge definition. We also evaluated the power to detect these effects if they were reduced by 15% and obtained 100% power with our sample size (CI: 99.63-100%).

#### 1.2 Experiment 2 – Perceived contrast for lighter and darker colours with sharp and blurred edges

**Best fitting linear mixed effect model for normalised contrast in the 2nd experiment (testing the effect of luminance pedestals and blurred edge)**, with fixed effects of size (2° - reference level, 0.5° and 0.33°), luminance pedestal (isoluminant – reference level, 3JND positive, 3 JND negative, 6 JND positive and 6 JND negative polarity), colour (reddish – reference level, greenish, bluish, yellowish) and edge (sharp – reference level, or blurred) and random intercepts only.

Model: normalised contrast ~ size+luminance pedestal+colour+edge+ size:luminance pedestal + luminance pedestal:edge + colour:luminance pedestal + colour:size + colour:edge + 1|participant

| **Fixed Effects** | **Estimate** | **t value** | **p value** | **95% CI** |
|---|---|---|---|---|
| Intercept | 109 | 363.852 | **<.001*** ** | 107-111 |
| 0.5 vs. 2 | 17.6 | 35.024 | **<.001*** ** | 14.5-20.7 |
| 0.33 vs. 2 | 24.8 | 46.449 | **<.001*** ** | 20.4-29.0 |
| 3 JND- vs. Isoluminant | -5.36 | -7.463 | **<.001*** ** | -8.01- -2.69 |
| 3 JND+ vs. Isoluminant | -1.89 | -2.669 | 0.008 | -3.98 – 0.181 |
| 6JND- vs. Isoluminant | -4.85 | -6.751 | **<.001*** ** | -7.60 - -2.06 |
| 6JND+ vs. Isoluminant | -2.84 | -3.901 | <.001*** | -5.67 – 0.107 |
| blur vs. sharp | -8.48 | -20.054 | **<.001*** ** | -10.4 - -6.62 |
| Green vs. Red | -2.30 | -3.804 | **<.001*** ** | -3.70 - - 0.909 |
| Blue vs. Red | 9.88 | 14.540 | **<.001*** ** | 7.39-12.5 |
| Yellow vs. Red | 3.30 | 5.153 | **<.001*** ** | 1.17-5.53 |
| 0.5 vs. 2 / 3JND- vs. Iso | 0.343 | 0.500 | 0.617 | -2.66 – 3.50 |
| 0.33 vs. 2 / 3JND- vs. Iso | -9.16 | -4.421 | **<.001*** ** | -14.1 - -4.10 |
| 0.5 vs. 2 / 3JND+ vs. Iso | 3.87 | 2.382 | 0.173 | 0.677 – 7.27 |
| 0.33 vs. 2 / 3JND+ vs. Iso | -5.12 | -2.590 | **0.0097** ** | -9.09 - -1.00 |
| 0.5 vs. 2 / 6JND- vs. Iso | 2.64 | 1.980 | 0.0478 | -2.49 – 8.04 |
| 0.33 vs. 2 / 6JND- vs. Iso | -8.26 | -3.755 | **<.001*** ** | -13.6 - -2.71 |
| 0.5 vs. 2 / 6JND+ vs. Iso | 4.31 | 2.750 | **0.0060** ** | 0.157 – 8.63 |

| | | | | |
|---|---|---|---|---|
| 0.33 vs. 2 / 6JND+ vs. Iso | -8.31 | -4.110 | **<.001*** ** | -12.7 - -3.71 |
| 3JND- vs. Iso / blur vs. sharp | -6.37 | -5.030 | **<.001*** ** | -9.13 – 3.64 |
| 3JND+ vs. Iso / blur vs. sharp | -1.81 | -1.272 | 0.203 | -4.58 – 0.962 |
| 6JND- vs. Iso/ blur vs. sharp | -6.78 | -5.253 | **<.001*** ** | -9.59 - -3.82 |
| 6JND+ vs. Iso / blur vs. sharp | -5.94 | -4.584 | **<.001*** ** | -9.50 - -2.45 |
| 0.5 vs. 2 / Green vs. Red | -2.84 | -1.851 | 0.064 | -5.54 - - 0.156 |
| 0.33 vs. 2 / Green vs. Red | -2.71 | -1.719 | 0.086 | -5.65 – 0.198 |
| 0.5 vs. 2 / Blue vs. Red | 11.9 | 7.642 | **<.001*** ** | 7.64 – 16.3 |
| 0.33 vs. 2 / Blue vs. Red | 16.4 | 9.686 | **<.001*** ** | 12.2 – 20.6 |
| 0.5 vs. 2 / Yellow vs. Red | 0.512 | 0.130 | 0.897 | -2.97 – 4.12 |
| 0.33 vs. 2 / Yellow vs. Red | 4.89 | 2.535 | **0.0113*** | 0.922 – 8.89 |
| 3JND- vs. Iso / Green vs. Red | -2.54 | -1.551 | 0.121 | -5.64 – 0.656 |
| 3JND+ vs. Iso / Green vs. Red | 0.769 | 0.337 | 0.736 | -1.96 – 3.48 |
| 6JND- vs. Iso / Green vs. Red | -1.17 | -0.639 | 0.523 | -4.22 – 1.86 |
| 6JND+ vs. Iso / Green vs. Red | -2.40 | -1.284 | 0.199 | -5.56 – 0.701 |
| 3JND- vs. Iso / Blue vs. Red | -2.55 | -1.218 | 0.223 | -7.13 – 2.30 |
| 3JND+ vs. Iso / Blue vs. Red | 7.16 | 3.408 | **<.001*** ** | 3.96 – 10.6 |
| 6JND- vs. Iso / Blue vs. Red | -5.54 | -2.635 | **0.0085**** | -9.57 - -1.30 |
| 6JND+ vs. Iso / Blue vs. Red | 0.527 | 0.250 | 0.802 | -3.71 – 5.02 |
| 3JND- vs. Iso / Yellow vs. Red | -6.86 | -3.261 | **0.0011**** | -10.6 - -3.13 |
| 3JND+ vs. Iso / Yellow vs. Red | -4.94 | -2.303 | **0.0214*** | -8.49 - -1.52 |
| 6JND- vs. Iso / Yellow vs. Red | -7.64 | -3.580 | **<.001*** ** | -11.3 - -4.01 |
| 6JND+ vs. Iso / Yellow vs. Red | -11.4 | -5.663 | **<.001*** ** | -14.8 - -8.19 |
| Blur vs. sharp / Green vs. Red | -2.79 | -2.522 | **0.0117*** | -4.49 - -1.04 |
| Blur vs. sharp / Blue vs. Red | -7.03 | -5.197 | **<.001*** ** | -9.12 - -4.97 |
| Blur vs. sharp / Yellow vs. Red | -2.75 | -2.075 | **0.0381*** | -4.54 - - 0.966 |
| *Random effects* | | | | |

| |
|---|
| Residual variance, $\sigma^2$ = 114.87 |
| Participant variance 42.73 |
| ICC 0.27 |
| N participant 20, 2273 Observations in total |
| Marginal $R^2$ / Conditional $R^2$ 0.537 / 0.663 |

The residuals for the initial model fit to the full dataset averaged across trials (2400 observations) were not normally distributed and suffered from outliers – removal of 127 outliers (i.e., those surpassing 1.96 SDs) was sufficient to correct the distribution of residuals, as evaluated by simulation-based residual diagnostics implemented in the DHARMa R package. To correct the distribution (which for a zero-bound variable such as contrast matching will be right-skewed), we fitted the model to log-transformed data and report the statistical results (F and t-values and p-values) from this model, while the estimates and 95% CIs are reported from the bootstrapped non-log-transformed model (with 1000 resamples of participants, as implemented in R package lmeresampler). Corresponding to the estimates, we report residual and participant variance from the non-log corrected model.

As evaluated by backward-reduced model fitting implemented in function *step* from R package lmerTest, The interactions that could be removed from the model were as follows: 4-way interaction (F(24,2253)=0.2635,p=1.00, all 3 way interactions (luminance pedestal by edge by colour: F(12,2253)=0.4705,p=0.933, luminance pedestal by size by edge: F(8,2253)=0.848,p=0.560), luminance pedestal by size by colour: F(24,2253)=0.988,p=0.478; size by edge by colour: F(6,2253)=1.303,p=0.252) and the 2-way interaction between size and edge (F(2,2253.1)=1.499,p=0.224). The following 2-way interactions could not be removed from the model: size by luminance pedestal (F(8,2253.1)=7.477,p<.001), luminance pedestal by edge (F(4,2253.1)=11.799,p<.001), size by colour (F(6,2253.1)=29.729,p<.001), luminance pedestal by colour (F(12,2253)=7.834,p<.001) and edge by colour (F(3,2253)=9.103,p<.001).

### POST-HOC ANALYSES FOR SIGNIFICANT 2-WAY INTERACTIONS

Post-hoc analyses were performed using *emmeans* package for R. We used the multivariate t distribution adjustment ('mvt') correction method for multiple comparisons for a series of t-tests across all the levels of the interaction.

#### *Interaction Between Factors of Colour and Size*

For the 2 degree stimulus, there were no significant differences between any of the colours (all ts <2.275, ps > .3697).

For the 0.5 degree stimulus, blue required significantly more contrast to achieve isosalience than red (OR 1.099 +/- 0.00863, t(2297) = 10.864, p <.001), green (OR 1.125 +/- 0.00837, t(2297) = 13.945, p <.001) and yellow (OR 1.090 +/- 0.0106, t(2297) = 8.825, p<.001).

Yellow also required more contrast to achieve equal salience than green (OR 1.046 +/- 0.00916, t(2297) = 4.944, p <.001) but not red (OR 1.018 +/- 0.00945, t(2297) = 1.888, p = .658). Finally, red also required more contrast for isosalience than green (OR 1.030 +/- 0.00974, t(2297) = 3.105, p = .0476).

For the 0.33 degree stimulus, a similar pattern was evident: blue required significantly more contrast to achieve isosalience than red (OR 1.125 +/- 0.00862, t(2297) = 13.512, p <.001), green (OR 1.149 +/- 0.00847, t(2297) = 16.194, p <.001) and yellow (OR 1.086 +/- 0.01093, t(2297) = 8.174, p<.001). Yellow also required more contrast to achieve equal salience this time against both green (OR 1.076 +/- 0.00927, t(2297) = 7.858, p<.001) and red (OR 1.050 +/- 0.00945, t(2297) = 5.109, p <.001). The weakly significant difference between red and green was this time below the significance threshold (OR 1.028 +/- 0.0101, t(2297) = 2.856, p = .0966).

***Interaction Between Factors of Size and Luminance pedestal***

For the 2 degree stimulus, there were no significant differences between different luminance pedestals (all ts <3.103, ps > .067).

For the 0.5 degree stimulus, the only significant differences involved the 3JND- stimulus, which required less contrast to achieve equal salience compared to both 3JND+ (OR 0.962 +/- 0.01032 SE, t(2297)=-3.638, p=.011) and 6JND+ (OR 0.958 +/- 0.01028 SE, t(2297)=-3.988, p=.003; all other ts < 2.536, ps > .289).

For the 0.33 degree stimulus, the isoluminant condition required more contrast to achieve equal salience in relation to all conditions with luminance pedestals:

3JND-: OR 1.098 +/- 0.0125 SE, t(2297) = 8.352, p<.001

3JND+: OR 1.056 +/- 0.0123 SE, t(2297) = 4.922, p<.001

6JND-: OR 1.095 +/- 0.0125 SE, t(2297) = 7.927, p<.001

6JND+: OR 1.083 +/- 0.0120 SE, t(2297) = 7.127, p<.001

Also, 3JND+ required significantly more contrast for isosalience compared to 3JND- (OR 1.039 +/- 0.0105 SE, t(2297) = 3.595, p=.0133) and 6JND- pedestals (OR 1.037 +/- 0.0115, t(2297) = 3.241, p = 0.0438).

***Interaction Between Factors of Colour and Luminance pedestal***

For red, the luminance pedestal did not affect perceived contrast (all ts < 2.256, all ps > .7895).

For green, the only significant difference was between isoluminant and 3ND-, the former requiring more contrast (OR 1.048 +/- 0.0130, t(2297) = 3.760, p = .0105; all other ts < 2.424, all ps > .4798).

For yellow, isoluminance required more contrast to achieve isosalience than all the luminance pedestals:

3JND-: OR 1.080 +/- 0.0138, t(2297) = 6.055, p<.001

3JND+: OR 1.065 +/- 0.0133, t(2297) = 5.019, p<.001

6JND-: OR 1.078 +/- 0.0138, t(2297) = 5.872, p<.001

6JND+: OR 1.098 +/- 0.0137, t(2297) = 7.485, p<.001

None of the other differences were significant ( all ts < 2.496, all ps > .429).

For blue, at isoluminance more contrast was required to achieve isosalience than for 6JND- luminance pedestal only:

3JND-: OR 1.042 +/- 0.0130, t(2297) = 3.289, p = .0554

3JND+: OR 0.964 +/- 0.0121, t(2297) = -2.924, p = .163

6JND-: OR 1.060 +/- 0.0134, t(2297) = 4.596, p<.001

6JND+: OR 0.991 +/- 0.0125, t(2297) = -0.742, p = 1.00

However, perceived contrast on positive luminance pedestals was reduced significantly when compared to those on negative pedestals:

3JND- vs. 3JND+: OR 0.925 +/- 0.0114, t(2297) = -6.136, p<.001

6JND- vs. 3JND+: OR 0.901 +/- 0.0137, t(2297) = -7.599, p<.001

3JND- vs. 6JND+: OR 0.951 +/- 0.0118, t(2297) = -4.061, p=.0035

6JND- vs. 6JND+: OR 0.935 +/- 0.0118, t(2297) = -5.363, p<.001

Differences between the two positive (OR 1.028 +/- 0.0128, t(2297) = 2.185, p = .678) or the two negative pedestals (OR 1.017 +/- 0.0126, t(2297) = 1.374, p = .995) were not significant.

***Interaction Between Factors of Edge Definition and Luminance pedestal***

For the sharp-edged stimulus, there were no significant differences between different luminance pedestals (all ts <2.219, ps > .3535).

For the blurred-edge stimulus, the isoluminant condition required more contrast to achieve equal salience in relation to all conditions with luminance pedestals except 3JND+:

3JND-: OR 1.080 +/- 0.00953, t(2297) = 8.762, p<.001

3JND+: OR 1.025 +/- 0.00899, t(2297) = 2.761, p = 0.105

6JND-: OR 1.078 +/- 0.00959, t(2297) = 8.391, p <.001

6JND+: OR 1.054 +/- 0.00936, t(2297) = 5.895, p<.001

The 3JND+ pedestal also required significantly more contrast to achieve isosalience with the 3JND- stimulus (OR 1.052 +/- 0.00830, t(2296) = 6.065, p <.001), the 6JND- stimulus (OR 1.052 +/- 0.00929, t(2297) = 5.714, p<.001) and the 6JND+ stimulus (OR 1.029 +/- 0.00906, t(2297) = 3.192, p = 0.0298). All other differences were non-significant (all ts < 2.920, all ps > .0896).

***Interaction Between Factors of Edge Definition and Colour***

The differences between different colours in terms of reduced perceived contrast through blurring of the standard stimulus edge was unequal: while highly significant, the effects were smaller for red:

Red: OR 0.947 +/- 0.00818, t(2297) = -6.595, p<.001

Green: OR 0.918 +/- 0.00845, t(2297) = -10.089, p<.001

Yellow: OR 0.928 +/- 0.00859, t(2297) = -9.295, p<.001

Blue: OR 0.866 +/- 0.00881, t(2297) = -13.703, p<.001

It is evident that the effect size for red is outside the 95% CI of the effect size for all other colours.

### SENSITIVITY ANALYSES FOR OBSERVED SIGNIFICANT FIXED EFFECTS

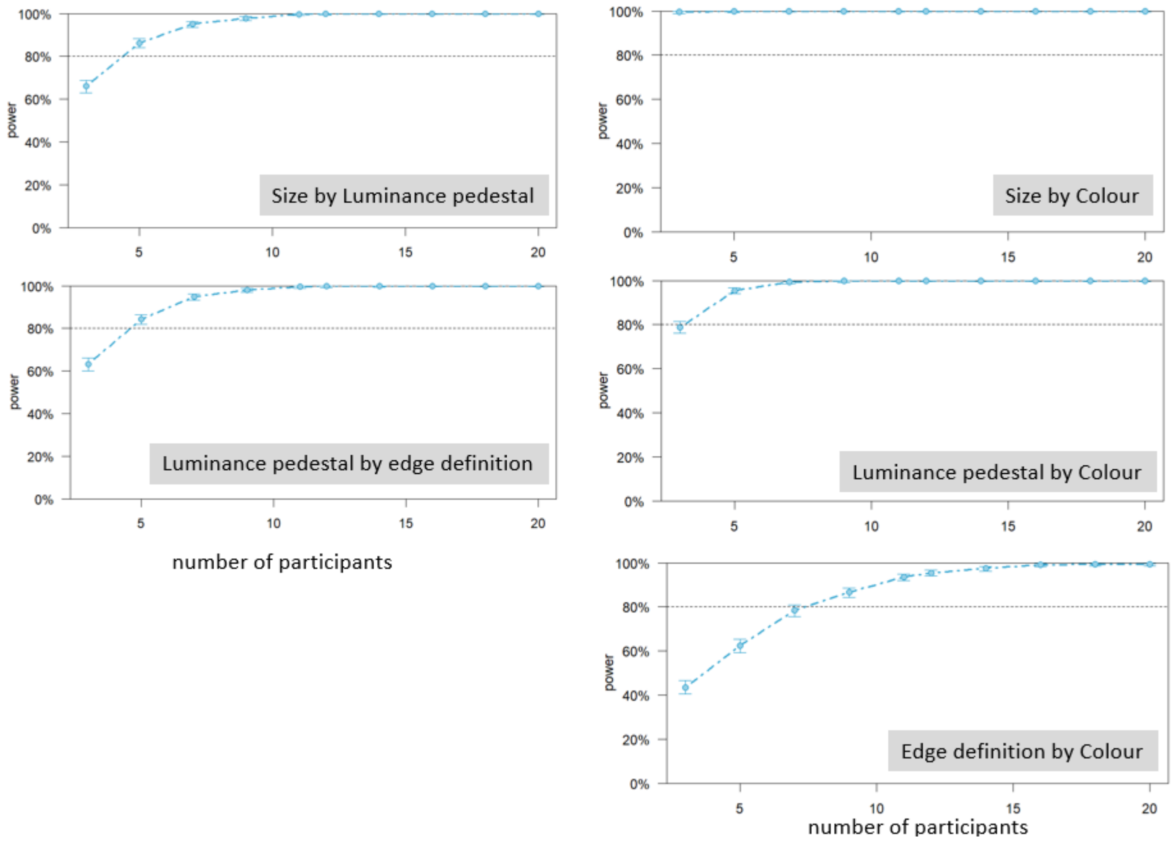

Sensitivity analyses were performed by resampling the number of participants (1000 simulations) and evaluating the observed power for obtaining the statistically significant fixed effects in the best fitting model, using the R package *simr*. Sufficient power for those effects can be observed with 8 out of 20 participants in all cases, with some effects being so strong as to require a very small number of participants to reach 80% power (indicated by a dotted line). Additionally, we reduced the effect sizes by 15% and evaluated the power to detect them, as recommended by Kumle and colleagues (Kumle et al., 2021). We had 100% power for detecting these reduced effects (CIs: 99.64-100%).

### VISUALISING DESATURATION EFFECTS

We have compiled a simulation of desaturation effects for isoluminant/equal contrast chromatic and achromatic sharp edge stimuli in Experiment 1. Disks are presented in the same size and their saturation adjusted to approximate the appearance of stimuli at smaller sizes. Effects below each disk describe the percentage of standard contrast in that condition, used to generate the disk. The background corresponds to the white point used in the experiment. Appearance of colours is not completely comparable to the experimental display.

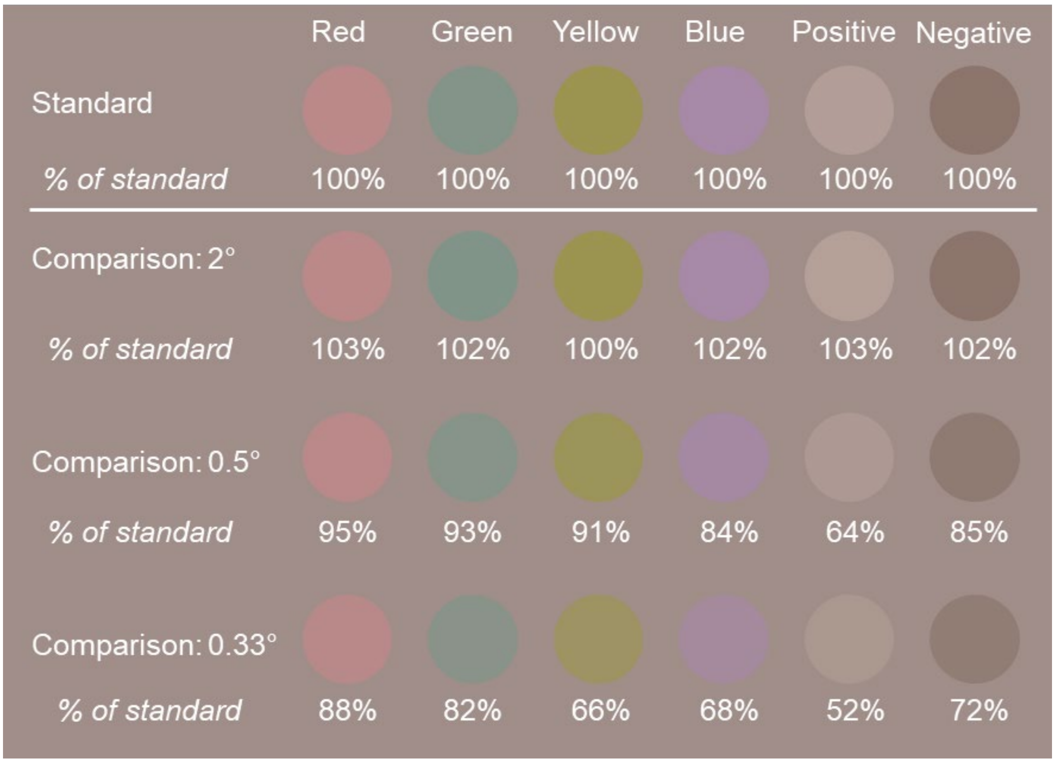
Red
Green
Yellow
Blue
Positive
Negative
Standard
% of standard
100%
100%
100%
100%
100%
100%
Comparison: 2°
% of standard
103%
102%
100%
102%
103%
102%
Comparison: 0.5°
% of standard
95%
93%
91%
84%
64%
85%
Comparison: 0.33°
% of standard
88%
82%
66%
68%
52%
72%
